# Concurrent multimodal imaging demonstrates that EEG-based excitation/inhibition balance reflects glutamate and GABA concentrations

**DOI:** 10.1101/2025.03.31.646339

**Authors:** Aaron Cochrane, Luke Rosedahl, Masako Tamaki, Takeo Watanabe, Yuka Sasaki

## Abstract

Neural excitation/inhibition (E/I) ratio is dynamically regulated on multiple timescales. Adaptive changes in E/I ratio can support healthy development, learning, and cognition, while disordered E/I ratio has been implicated in neurodevelopmental disorders, neurodegenerative disorders, and states of impaired vigilance. There has been growing interest in inferring E/I ratio from efficient and noninvasive measurements such as electroencephalography (EEG), and several algorithms have been proposed to estimate E/I ratio from EEG. Despite promising results, there has been a lack of validation studies testing the underlying neurochemical changes leading to increased or decreased EEG-based E/I ratio. Here, using concurrent EEG and magnetic resonance spectroscopy (MRS) for over an hour, we assessed which algorithm of EEG-based E/I ratio best matched with MRS-based E/I ratio in humans of both sexes. The MRS-based E/I ratio was obtained by the ratio of glutamate concentration to GABA concentration. We applied 10 candidate indices of EEG-based E/I ratio using four approaches in several spontaneous frequency bands. Uniquely, we quantified the associations between the EEG-based E/I ratio and MRS-based E/I ratio separately for between-subjects and within-subjects variations. We found that each EEG-based E/I algorithm showed reliable and positive associations with MRS-based E/I, and especially EEG-based E/I ratio in alpha band with a criticality theory based approach showed the best association to the MRS-based E/I ratio. While these associations were evident for between-subjects comparisons, they were quite weak for within-subjects comparisons. These results suggest that EEG-based E/I algorithms are likely to reflect, at least in part, relative concentrations of glutamate and GABA.

## Introduction

Brain functioning is largely regulated through a dynamically changing balance of excitatory and inhibitory (E/I) activities (Hensch et al. 1998; Shew et al. 2011). Such dynamics are closely linked to a variety of phenomena including neural plasticity (Bavelier et al. 2010; Hensch et al. 1998; Vogels et al. 2011), brain development (Hensch 2004; Takesian and Hensch 2013) disordered functioning (Bruining et al. 2020; Geertjens et al. 2022; Nifterick et al. 2023), and cognition-related phenomena (Hellyer et al. 2016; Yang et al. 2023; Amil and Verschure 2021). Reflecting the fact that E/I ratio varies on multiple spatial and temporal scales, and is difficult to measure non-invasively (Ahmad et al. 2022; Craven et al. 2022), a variety of methods are used to assess E/I in humans. These include the use of electrophysiological (EEG) signatures and magnetic resonance spectroscopy (MRS). Each of these approaches holds great promise and yet faces serious limitations.

EEG-based approaches to quantifying E/I ratio (i.e., EEG-E/I) come from several theoretical and computational sources, including scale-free dynamics (Gao, Peterson, and Voytek 2017), criticality theory (Poil et al. 2012), and neural mass modeling (Yokoyama and Kitajo 2023), and have been used to investigate physiological underpinnings of phenomena ranging from vigilance (Yang et al. 2023) to Alzheimer’s disease (Nifterick et al. 2023). Each of these has proposed mechanistic bases for the links between a scalp-based electrical signal and cortical excitation or inhibition, but empirical data tracking brain regions’ neurochemistry to EEG measures has been largely lacking (Ahmad et al. 2022; Geertjens et al. 2022). In contrast, EEG-based E/I ratio measures have relative advantages include their wide availability, inexpensive and efficient implementation, and providing data from multiple brain regions simultaneously.

The use of MRS to study E/I ratio (i.e., MRS-E/I) ratio presents a different and complementary set of trade-offs. Studies of E/I ratio using MRS have been conducted to study brain plasticity, including that involved in perceptual learning (Shibata et al., 2017; Tamaki et al., 2020). While MRS-based E/I has the distinct benefit of having a clear biological basis, namely, excitatory (glutamate) and inhibitory (gamma-aminobutyric acid; GABA) neurotransmitter concentrations, it is limited to a single cortical location with fairly low temporal resolution and restricts the measurement only inside an MRI scanner, thereby precluding measurement in certain populations or during certain behaviors, and is relatively expensive.

There have been efforts to link EEG measures and MRS data. There have been multiple reports of associations between MRS-measured GABA concentrations with electrophysiological frequency bands (e.g., Balz et al., 2016; Edden et al., 2009), but each of these studies measured electrophysiology and MRS in separate sessions and were therefore limited to stable inter-individual correlations and were insensitive to rapid or functional changes that may occur on short timescales. However, functional changes in varying E/I ratio would be necessary to investigate in many fields, including neuroplasticity, pharmacological interventions, and cognition.

There remains a clear need for multimodal investigations of EEG-E/I ratio in conjunction with MRS-E/I ratio. Only simultaneously measuring the two forms of E/I ratio can allow assessment of both within-subject and between-subject variations in each of the two measures. For example, it is possible that the two forms of E/I ratio are empirically distinct from each other, if EEG-based E/I ratio reflects specific neural subpopulations generating oscillatory signatures (Scheeringa and Fries 2019; Shafiei et al. 2023), while MRS-based EI ratio reflects the aggregate concentration of certain molecules in a large voxel. In contrast, it is possible that the two forms of E/I are indexing the same underlying biological processes, that is, the relative concentrations of the primary excitatory vs. inhibitory neurotransmitters in a broad region of cortex. Here we addressed these questions by comparing MRS-based E/I ratio with EEG-based E/I ratio using several approaches in participants in simultaneous measurements of MRS and EEG in healthy young participants.

## Materials and Methods

### Experimental Design

We re-analyzed data from Study 1 of Tamaki et al. (2020), which consists of 18 subjects who slept for between 60 and 90 minutes following a perceptual learning task. Simultaneous polysomnography including EEG and MRS were recorded throughout the sleep session. To assess the extent to which candidate EEG-based measures of E/I ratio were reflecting the changes in neurotransmitter concentrations measured by MRS-based E/I ratio, we estimated a single E/I value using each method for each 5-minute period of the recorded data (see sections *MRS-based E/I ratio* and *EEG-based measures of E/I ratio*). We then used Bayesian multivariate linear mixed-effects modeling and cross-validation to quantify the ability to predict MRS-based E/I using EEG-based estimates (see section *Statistical Analysis*). We separated these estimates into between-subjects and within-subjects analyses; while much previous work has focused on the relatively stable differences between individuals’ levels of excitation and inhibition (e.g., Balz et al., 2016; Edden et al., 2009), within-subjects variation in E/I ratio are central to many research domains such as vigilance (Yang et al. 2023) and learning-related neuroplasticity (Bavelier et al. 2010; Tamaki et al. 2020) because these topics are inherently concerned with intra-individual changes. Separating these results helps identify both the overall associations between MRS-E/I ratio and EEG-E/I ratio as well as their relative isomorphism in the context of different applications.

### MRS-based E/I ratio

MRS-based E/I ratio is typically estimated by identifying a brain region of interest (e.g., early visual areas) and placing a single voxel in that region. Using magnetic resonance, the spectroscopic signal is then measured in that voxel over the course of several minutes and the relative concentrations of excitatory and inhibitory neurotransmitters are estimated (Provencher, 1993). These concentrations can then be examined separately or combined into a single measure of E/I ratio. Using such methods with a 3 Tesla MRI it would conventionally be considered high spatial resolution to have a single voxel that is 15mm per side and a high temporal resolution to have one measurement every 2 minutes. The relative trade-offs of MRS measures of E/I ratio are clear, then: While MRS-based E/I ratio is a relatively direct measure of actual molecules of interest within the brain, this measurement technique is resource-intensive, spatially and temporally imprecise, and the measured spectrum is itself a noisy measure. Indeed, the estimation of neurotransmitter concentrations has no consensus standard and results from a single MR sequence may vary depending on the estimation algorithm (e.g., inter-algorithm correlations between GABA concentration estimates ranging from below .1 to above .8; Craven et al., 2022).

Our primary results used the originally-published MRS data (Tamaki et al. 2020; Study 1), wherein a MEGA-PRESS sequence was used (1.25s TR; 2.2 × 2.2 × 2.2 cm voxel, placed on midline in early visual cortices; 3 Tesla Siemens scanner). We made two modifications to the data reported in the previous publication, namely, partial volume correction (PVC) to account for relative proportions of grey matter (i.e., to mitigate the influence of grey matter proportion on associations with EEG measures) and separation into 5-minute sections of time instead of 10-minute time sections (mean number of 5-minute windows per participant = 13.1, sd = 3.6). As with the originally-published work, we used LCModel software (Provencher, 1993) to identify Glx (Glutamate + Glutamine) and GABA+ (GABA + residual molecules) concentrations normalized to NAA. We then estimate MRS-based E/I ratio by dividing Glx by GABA+. We chose to continue using LCModel to estimate Glx and GABA+, rather than other candidate softwares for several reasons. First, we needed to maintain congruence with the original findings because reliable results in the original paper indicate that E/I from LCModel reflects true underlying cortical changes (Tamaki et al., 2020). Second, LCModel has been found to provide the most typical estimates of neurotransmitter concentrations, as it shows the highest correlation of GABA estimates with the median of estimates from other software (Craven et al., 2022).

### EEG-based measures of E/I ratio

We assessed four approaches to quantifying electrophysiological activity from EEG. Three of these approaches were calculated separately for three canonical frequency bands (see below), leading to a total of 10 total EEG-based measures to test associations with MRS-E/I. The first measure, overall amplitude, is not a measure of E/I *per se*, but associations between MRS-E/I within specific frequency bands have been observed previously (e.g., theta and low beta; Tamaki et al. 2020). Next, we assessed EEG-based E/I ratio derived from scale-free dynamics reflected by the 1/f power slope (Gao, Peterson, and Voytek 2017), then an extension of this scale-free perspective that derives from criticality theory (Bruining et al. 2020), and a set of estimates utilizing a neural mass model of excitatory and inhibitory dynamics (Yokoyama and Kitajo 2023).

For details of EEG preprocessing steps, such as MR artifact rejection, see Methods in Tamaki et al. (2020; Study 1). In all analyses we report EEG measures estimated from the closest 7 electrodes (Oz, O1, O2, PO3, PO4, & PO7) to the MRS voxel, since the MRS voxel was placed over early visual areas (Tamaki et al., 2020). We show the detailed information for each approach below.

#### Frequency band amplitude

As one of the most basic measures identifiable within EEG signals, we included simple amplitude (i.e., square root of power) measures from several frequency bands. We used finite impulse response (FIR) bandpass filtering in EEGLAB (Delorme & Makeig, 2004) to isolate theta (5-8Hz), alpha (8-12Hz), and low beta (12-20Hz) bands, estimated the amplitude in 5-second windows, and then calculated the trimmed mean amplitude for each 5-minute measurement period (i.e., for robustness to outliers, after removing the 5% lowest and 5% highest amplitude windows, calculating the arithmetic mean of the amplitudes within the remaining windows). These three frequency bands were examined due to previous work finding links between MRS-based E/I ratio and theta as well as low beta power (Tamaki et al., 2020), and the use of alpha amplitude envelopes in the implementation of the criticality-based E/I ratio (Bruining et al., 2020).

While there have been some indications of links between frequency bands and E/I ratio (Balz et al., 2016; Duncan et al., 2014; Porjesz et al., 2002), we include amplitude here primarily as a baseline approach against which to assess the more algorithmically complex approaches to estimating EEG-based E/I ratio. That is, as an algorithmically simple derivation from resting-state EEG, we consider amplitudes to be valuable measures to assess overall data reliability (e.g., see Supplement *Across-channel correlations*; we would expect amplitude measures to have high cross-channel correlations and a lack of such correlations would indicate poor data quality), test the divergence of EEG-E/I measures from amplitudes (see Supplement *Associations among EEG measures*), as well as providing parsimonious reflections of MRS measures if the relevant correlations were to be found (Edden et al., 2009; Tamaki et al., 2020).

#### EEG-E/I from 1/f slope (“1/f”)

The next approach to derives E/I ratio from EEG signals is based on the widespread observation that neural activity, in temporal, spatial, and frequency domains, displays scale-free dynamics (power-law, or “1/f” when in the frequency domain; Hahn et al., 2017; He, 2014; Linkenkaer-Hansen et al., 2001). Computational derivations of the slopes of these 1/f distributions have been linked to cortical E/I ratio due to the relative timescales of excitatory and inhibitory interneurons’ firing rates (Gao et al., 2017; Lendner et al., 2020). As with previous work (Gao, Peterson, and Voytek 2017; Yokoyama and Kitajo 2023) we estimated the 1/f slope coefficient between 30Hz and 50Hz (bandpass filtered) for each 5-minute window of interest, which avoids contamination of the 1/f slope with deviations from this slope associated with alpha and beta frequencies; we implemented this approach with Python scripts available in conjunction with Yokoyama and Kitajo (2023). Unlike the other approaches, 1/f slope does not consider discrete frequency bands, but instead the change in power across a wide range of frequencies. The following approach, based in criticality theory, is an elaboration of these principles of scale-free dynamics.

#### EEG-E/I from criticality theory (“fE/I”)

A recently-developed algorithm for identifying E/I ratio from EEG data is grounded in the computational biology framework of criticality, which characterizes the nature of many biological systems as existing at the cusp of state transitions (Bak 1999; Márton et al. 2014; Poil et al. 2012). One application of this theoretical approach is to neural criticality, which proposes that neural activity exists approximately at a ratio between excitation and inhibition in order to maximize the flexibility of macro-scale neural computations (Avramiea et al. 2022; Cocchi et al. 2017; Shew et al. 2011). To estimate this type of EEG-based E/I ratio (Ahmad et al., 2022; Juarez Martinez et al., 2019), we used an algorithm which is based in criticality theory and has been validated in computational, pharmacological (Bruining et al. 2020), and animal models (Kat et al., 2024); the developers of this algorithm label it “functional E/I ratio” (fE/I). We do not know of any previous work linking fE/I to MRS measures.

We applied this approach to the same three bandpass-filtered frequency bands we considered when examining simple amplitude measures. In brief, this algorithm first determines the degree of *criticality* (here, linked to the exponent of Detrended Fluctuation Analysis (DFA) Long-Range Temporal Correlations (LRTC)). This uses links between the window size of an analyzed signal and the variability in its cumulative detrended amplitude profile. Second, this degree of criticality is linked to the signal amplitude; in inhibitory-dominant regimes these two values should be positively correlated, while in excitation-dominant regimes they should be anticorrelated (Bruining et al., 2020). Using code published in conjunction with Bruining et al. (2020), fE/I values were estimated for each of the above-defined three frequency bands (theta, alpha, and low beta) and then within- and between-subject correlations with MRS-E/I were estimated. As in the publication establishing the method, estimation of fE/I used a window size of 5 seconds with an inter-window overlap of .8 (Bruining et al. 2020); we also tested the effects of increasing the window sizes to 15 seconds and found minimal changes, and here report only 5-second window sizes for simplicity. For each 5-minute measurement period we also estimated the DFA coefficient and excluded measurement periods with DFA below .6, as recommended by the developers of this approach (Bruining et al. 2020; Diachenko et al. 2024).

#### EEG-E/I from neural mass model (“NMM-E/I”)

A fourth approach comes from a body of work instantiating neural mass models, that is, computational models of neural circuitry such as cortical columns; such models can be scaled up and can simulate complex dynamics and characterize dynamic and nonlinear features of neural activity (Cooray, Rosch, and Friston 2023; Jansen and Rit 1995). Yokoyama and Kitajo (2023) extended earlier neural mass models (Jansen and Rit 1995), which had been developed for purposes such as modeling epileptic dynamics, with a key insight: such models can include estimates of the relative activity of circuits’ excitatory and inhibitory neural populations. Following this, and so they could compare to the 1/f approach described above, Yokoyama and Kitajo (2023) showed that the E/I ratio derived from such models could capture the fluctuating activity of humans’ electrophysiological activity during sleep. This approach to estimating E/I ratio is quite new, however, and to our knowledge has not been widely assessed. We estimated this NMM-E/I using Python code available in conjunction with Yokoyama and Kitajo (2023) for each 5-minute measurement period, and for each of the canonical frequency bands described above (i.e., theta, alpha, & low beta). Unlike the fE/I algorithm, the NMM-E/I ratio algorithm does not involve separation into 5-second segments. We also tested the NMM-E/I on a single wider set of frequencies, more similar to the approach implemented in Yokoyama and Kitajo (2023), on bandpass-filtered data between 5Hz and 20Hz (i.e., the lower and upper limits of the three frequency bands considered in the above analyses). The resulting NMM-E/I values did not improve correspondence to MRS-E/I relative to the values estimated from distinct canonical frequency bands.

### Statistical Analysis

For both within-subjects and between-subjects analyses we used two separate statistical approaches which were complementary to one another. In the first approach, in order to best estimate the effect size (i.e., correlation coefficient) between each EEG measure and MRS-based E/I ratio, we fit a Bayesian multivariate linear mixed-effects model using **brms** in R (Bürkner 2017). Using this approach we specified each of the measures as variables to predict (i.e., “response” variables). Each response variable was fit with an overall intercept, a random-effects by-participant intercept, and a fixed-effect autoregressive term with an order of one to account for non-independence of temporally proximal measurements. Within the multivariate model the correlations across the response variables of the by-participant intercepts were estimated (in brms “Wilkensen” notation the formula for each E/I measure was `response ∼ 1 + ar(time, order = 1) + (1 | subjectR | subjectID)`; for full schematic see Figure S1). In this way we estimated simultaneously the between-subjects effects, in the form of the correlations between different response variables’ by-participant intercepts, as well as within-subjects effects, in the form of the residual correlations remaining after accounting for the autoregressive and by-subject terms. We report these two correlations, both between-subjects and within-subjects, as the best empirical estimate of the links between each measure. In addition, this model included an estimation of the remainder of the correlation matrix (e.g., between all EEG-E/I measures as well as estimates of Glx and GABA+); we report these correlations to determine the extent to which our EEG measures are isomorphic and are potentially more reflective of Glx or GABA+ variations (see Figure S2).

As tests of the reliability of associations between EEG measures and MRS-based E/I ratio, and in consideration of the possibility of overfitting our data, we augmented our Bayesian multivariate linear mixed-effects model with a maximum-likelihood cross-validation approach. This approach involved iteratively and randomly partitioning the data into “training” and “test” sets, fitting a maximum-likelihood linear regression model predicting MRS-based E/I ratio to the training set, and assessing the R-squared (i.e., proportional reduction in sum of squared error; out-of-sample delta R-squared dRsq_oos_) on the test set (Dale et al., 2021; Scott et al., 2023). Between-subjects and within-subjects analyses were necessarily implemented somewhat differently. Between-subjects cross-validation involved first calculating the subject-level means for each measure, and then using simple random partitioning of this aggregated dataset into 80%/20% subsets, with data being fit with a bivariate robust linear model to 80% of the data and dRsq_oos_ assessed on the remaining 20%. The null model in this case was an intercept-only linear regression. In contrast, within-subjects cross-validation involved randomly selecting a single observation (i.e., 5-minute time period) from each participant to keep as a test set, fitting a linear model including a by-participant covariate (i.e., to control for participant-level means in MRS-E/I ratio) to the remaining training set, and then assessing the proportional change in out-of-sample squared error (dRsq_oos_) when compared to a model with only the participant covariate as a predictor. To control for between-subjects variations in the predictor EEG-based measures, for each of these within-subjects cross-validation models the EEG-E/I measure was first normalized within participants, after using a Yeo-Johnson transformation to minimize univariate skew, and then z-scored. Here we report the median dRsq_oos_ of 500 resampled models for each EEG-E/I measure.

### Supplemental Analyses

#### EEG-E/I reliability assessment

We implemented a cross-channel correlation approach to estimate a heuristic reliability of our EEG data and EEG-based measures. Due to basic principles of volume conduction (Burle et al., 2015), as well as our intention to measure relatively large-scale variations in physiological activity, we expected highly homogeneous estimates of EEG measures across our measured channels. A lack of such a correlation for all EEG measures would indicate poor EEG data quality, whereas a lack of correlation within a single EEG measure may indicate computational instability of that measure. Because each of our EEG-based methods involved first estimating a value for each channel we were able to, within each method, estimate the cross-channel correlations for our analyzed posterior electrodes. We tested these correlations within subjects by finding the pairwise cross-channel partial rank correlations in estimated E/I values, partialling out participant-level intercepts. We then report the median cross-channel partial rank correlation (see Figure S3). While we do not have a strict reliability criterion for these correlation measures, we believe that the relative cross-channel correlations should provide a general sense of the computational stability of the measures; amplitude, fE/I, and 1/f measures demonstrated acceptable reliability.

#### Between-subjects variability relative to within-subjects variability

In light of differences in between-subjects relative to within-subjects correspondences between EEG-based and MRS-based measures of E/I, we sought to quantify the relative between-subjects and within-subjects variances for each measure. To determine between-subjects variances we first found each participant’s median value for each measure, then calculated the variances of these median values. To determine within-subjects variances we first calculated each participant’s variance for each measure, then found the median of these variances. The ratio of between-subjects variance to within-subjects variances is reported in Figure S4, with no clear bias toward larger between-subjects variance nor within-subjects variance.

## Results

We first assessed data quality and excluded 5-minute windows in which at either fE/I ratio or MRS-E/I ratio demonstrated high noise relative to signal; amplitude, 1/f, and NMM-E/I ratio measures do not have established cutoff thresholds for exclusion. MRS data quality was quantified using the ratio of the uncertainty of molecules’ estimated concentration relative to the corresponding estimate; as in the originally published work (Tamaki et al. 2020), a 20% level of the Cramer-Rao lower bound was used for MRS exclusion. We assessed EEG quality using the detrended fluctuation analysis (DFA) exponent coefficient criterion which Bruining et al. (2020) describe in conjunction with the fE/I method; this involves estimating the DFA coefficient and excluding windows in which the coefficient is less than .6. While the original sample included 18 participants’ datasets, three participants had only three or fewer remaining MRS values and were excluded from further analysis. This left 15 participants data, with a mean number of 5-minute samples of 13.1 (i.e., 65.5 minutes).

### Between-subjects associations between EEG measures and MRS-based E/I

Inter-individual differences in certain measures of EEG-E/I were correlated with MRS-E/I (see Figure 1 & Table 1). Each of the candidate algorithms produced E/I estimates with more than 5% of cross-validated predictive variance explained in at least one frequency band (i.e., dRsq_oos_ > .05); we consider each of the links with positive dRsq_oos_ values to be reliable. The criticality-based fE/I approach derived from alpha oscillatory activity was most closely linked to MRS-E/I across participants, followed by 1/f slope, then NMM-E/I derived from low beta activity, and then fE/I derived from theta band. All other measures, including overall amplitudes, were not reliably predictive of inter-individual differences in MRS-E/I. The lack of association with overall amplitudes provides a justification for the candidate algorithms’ approaches to estimating E/I ratio; further, the strong performance of criticality-based alpha-band E/I supports the developer’s findings regarding this algorithm (Bruining et al. 2020; Diachenko et al. 2024).

**Figure 1.**
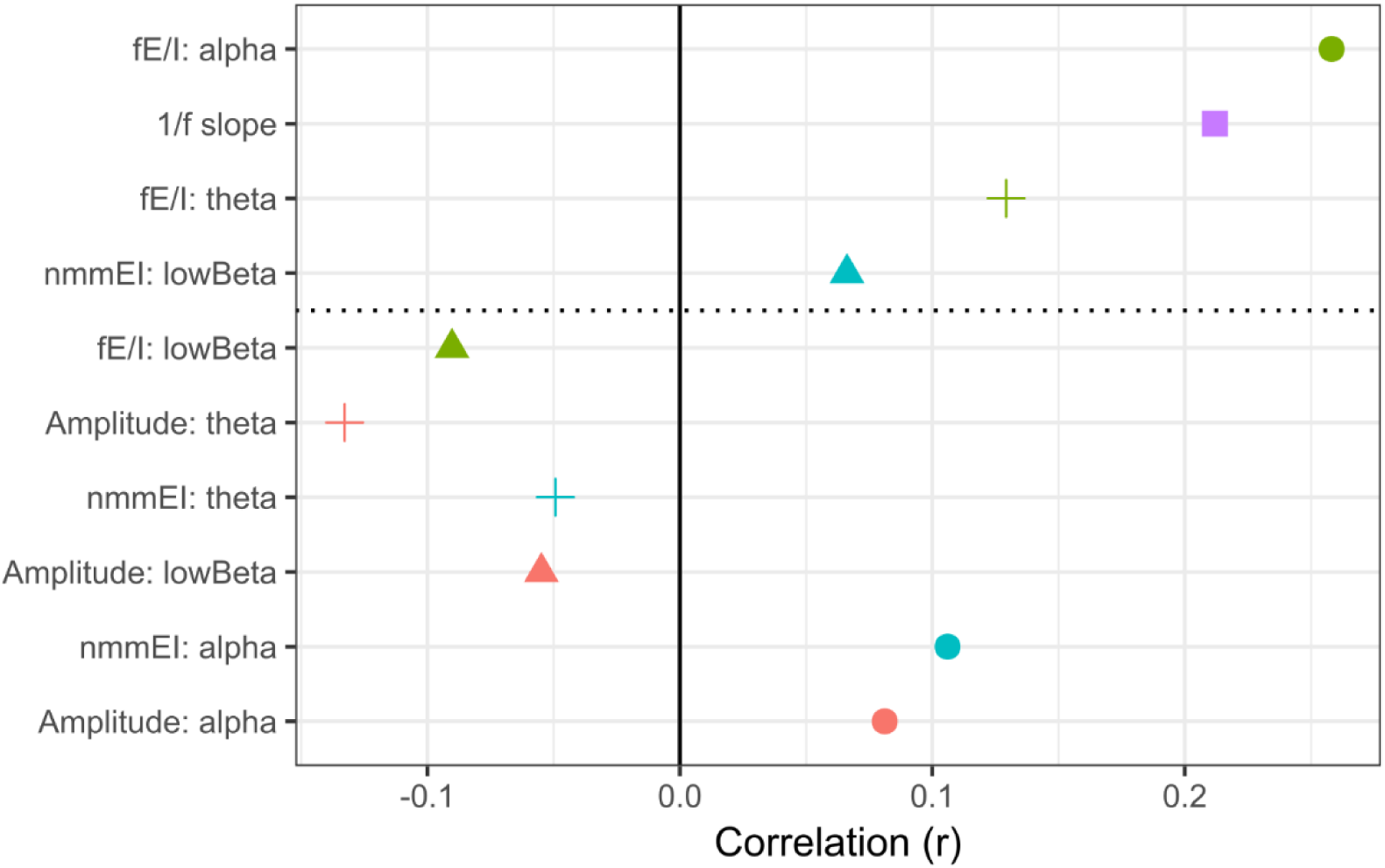
Between-subject associations between EEG-E/I measures and MRS-E/I. Correlation of by-participant random intercepts from the Bayesian multivariate mixed-effects model Each panel is sorted by decreasing dRsq_oos_ (i.e., measures at the top have the highest out-of-sample variance explained). Dashed line separates variables explaining reliable out-of-sample variance with those that do not; we consider all associations above the dashed line to be reliable.

**Table 1.** Between-subject associations between EEG-E/I measures and MRS-E/I. Each EEG measure’s association with MRS-based E/I ratio is shown in the form of cross-validated out-of-sample variance explained (dRsq_oos_) and effect size extracted from a Bayesian multivariate mixed-effects model (r). All dRsq_oos_ values above zero are considered to be reliable, with values below replaced by “negative”.

| EEG measure | $dRsq_{oos}$ | Correlation (r) |
| --- | --- | --- |
| 1/f slope | 0.131 | 0.211 |
| Amplitude: alpha | negative | 0.081 |
| Amplitude: lowBeta | negative | -0.054 |
| Amplitude: theta | negative | -0.132 |
| fE/I: alpha | 0.190 | 0.258 |
| fE/I: lowBeta | negative | -0.09 |
| fE/I: theta | 0.112 | 0.129 |
| nmmEI: alpha | negative | 0.106 |
| nmmEI: lowBeta | 0.103 | 0.066 |
| nmmEI: theta | negative | -0.049 |

### Within-subjects associations between EEG measures and MRS-based E/I

After controlling for participant-level effects, several approaches to EEG-E/I were reliably associated with MRS-E/I (i.e., dRsq_oos_ > 0; see Figure 2 & Table 2). While this wide range of EEG-E/I approaches providing within-subjects out-of-sample predictiveness for MRS-E/I is promising, a closer investigation of the effects tempers this promise somewhat. First, although out-of-sample variance explained was positive, it was quite small, with only simple amplitude measures providing more than 1% of predictive value. Second, of the EEG measures with positive dRsq_oos_ values, only theta amplitude had a positive correlation with MRS-based E/I ratio (r =.08). These results are in striking contrast to the between-subjects findings, where a relatively large amount of variance was explained with positive relations between EEG-E/I and MRS-E/I measures.

**Figure 2.**
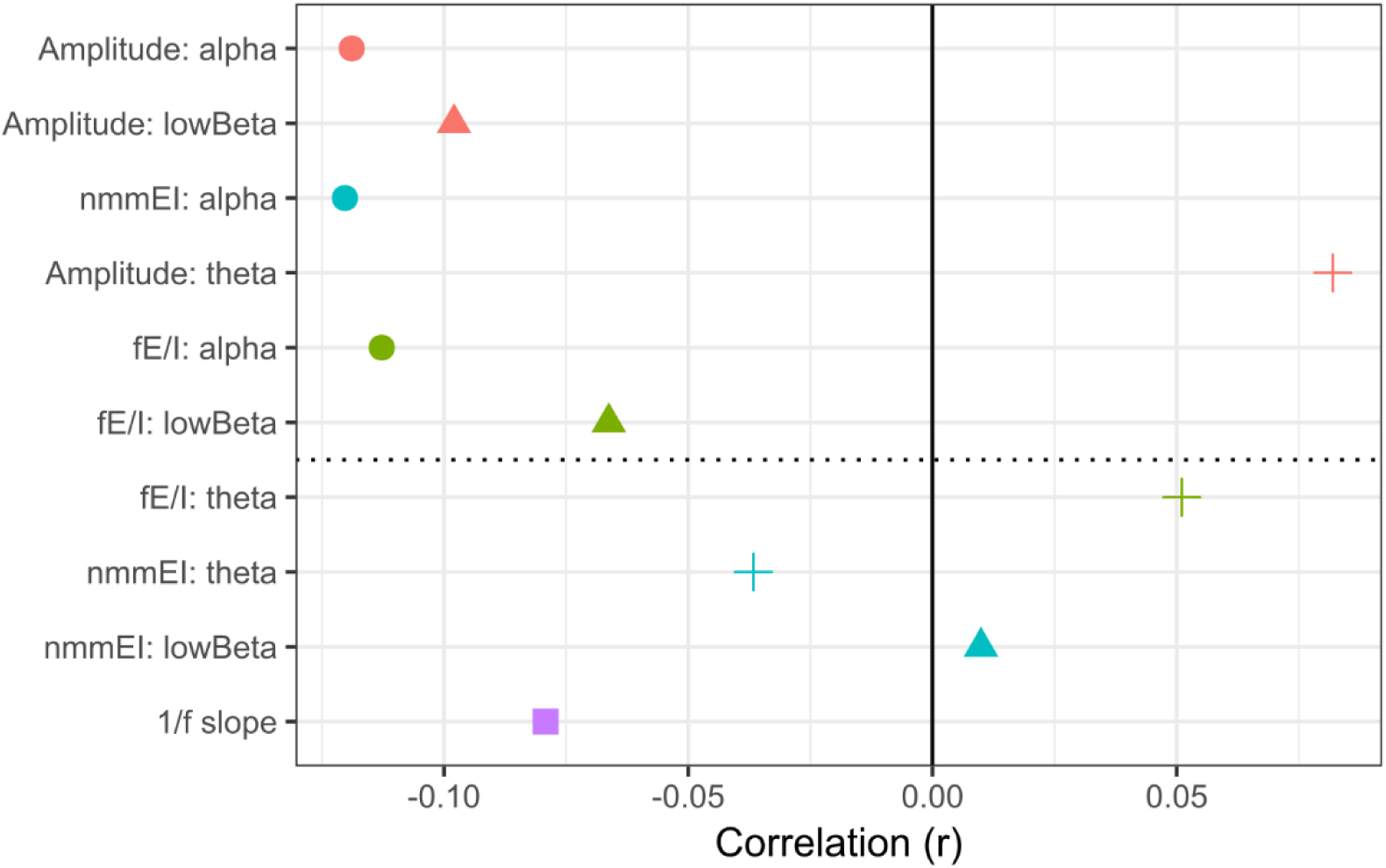
Within-subject associations between EEG-E/I measures and MRS-E/I.) Correlation of by-participant random intercepts from the Bayesian multivariate mixed-effects model. Each panel is sorted by decreasing dRsq_oos_ (i.e., measures at the top have the highest out-of-sample variance explained). Dashed line indicates whether variables explained out-of-sample variance or did not; we consider all associations above the dashed line to be reliable.

**Table 2.** Within-subject associations between EEG-E/I measures and MRS-E/I. Each EEG measure’s association with MRS-based E/I ratio is shown in the form of cross-validated out-of-sample variance explained (dRsq_oos_) and effect size extracted from a Bayesian multivariate mixed-effects model (r). All dRsq_oos_ values above zero are considered to be reliable, with values below replaced by “negative”.

| EEG measure | $dRsq_{oos}$ | Correlation (r) |
| --- | --- | --- |
| 1/f slope | negative | -0.079 |
| Amplitude: alpha | 0.021 | -0.118 |
| Amplitude: lowBeta | 0.013 | -0.097 |
| Amplitude: theta | 0.005 | 0.081 |
| fE/I: alpha | 0.004 | -0.112 |
| *fE/I: lowBeta* | 0.001 | -0.066 |
| *fE/I: theta* | negative | 0.051 |
| *nmmEI: alpha* | 0.009 | -0.120 |
| *nmmEI: lowBeta* | negative | 0.009 |
| *nmmEI: theta* | negative | -0.036 |

## Discussion

We have seen that EEG can be used to estimate E/I ratio in a manner that is compatible with, but not wholly overlapping with, variations in E/I ratio estimated from MRS. This mechanistic basis in the ratio between glutamate and GABA was true for several distinct approaches to EEG-based E/I ratio. In particular, EEG-based E/I ratio between-subjects was associated with MRS-based E/I ratio between-subjects, to the extent that they are comparable to (or even larger than) the correlations between MRS neurotransmitter concentrations found when comparing different estimation algorithms for MRS measures of GABA (Craven et al. 2022). In particular, the “functional E/I” algorithm derived from criticality theory was most closely associated with between-subjects differences in MRS-based E/I ratio; this reliance on temporal variations in alpha power corroborates the work of the developers of the fE/I algorithm (Bruining et al., 2020; Diachenko et al., 2024). Beyond this criticality-based EEG-based E/I ratio measure, each of the other EEG-based E/I ratio approaches were also positively associated with between-subjects variations in MRS-based E/I ratio. These results showed broad support for each of the approaches, which were themselves not highly correlated (see Figure S2). A lack of isomorphism between the EEG-based measures means that future work may benefit from combining these algorithms to best characterize cortical E/I. The relative contribution of estimates from separate frequencies is of particular interest; the generative mechanisms underlying variations in theta, alpha, and beta rhythms provides the eletrophysiologist with a rich and multidimensional signal from which to infer shifts in cortical states. Indeed, independent frequency bands have been previously associated with glutamate and/or GABA levels (Porjesz et al., 2002; Tamaki et al., 2020), and a broad spectrum of frequencies showed links to MRS-based E/I ratio when analyzed with different algorithms in the current work (i.e., theta and alpha analyzed with fE/I; low beta analyzed with a neural mass model; gamma when calculating a 1/f slope). We believe that combining amplitude and E/I estimates from several frequency bands may provide additional precision, but we are hesitant to attempt such a combinatorial search within our limited dataset.

In contrast to between-subjects patterns of E/I ratio, within-subjects associations between MRS-based E/I ratio and EEG-based E/I ratio were much weaker; indeed, only simple low beta amplitude was reliably associated with within-subjects variations in MRS-based E/I ratio in the same direction (i.e., negative) as with between-subjects variations in MRS-based E/I ratio. The reasons for the weakness of these associations cannot be known with certainty, but it is likely to arise partly to a decrease in signal strength and an increase in noise when compared to between-subject comparisons. That is, estimating EEG-based E/I ratio as well as MRS-based E/I ratio on windows of only 5 minutes is likely to lead to relatively noisy estimates; while shorter measurement windows measuring GABA with MRS have been used in the past (Frank et al. 2022), longer windows (e.g., 10 minutes) are more standard. Unfortunately our available data is limited enough that estimating such within-subjects associations between EEG-based E/I ratio and MRS-based E/I ratio appears to be largely overwhelmed by the reduced signal-to-noise ratio; the MR context of these estimates may also have itself reduced data quality in estimating EEG-based E/I.

There may be alternative sources of divergences between EEG-based and MRS-based E/I ratios. On one hand, the neural mechanisms giving rise to each of our measured signals may be distinct; while MRS measured bulk concentrations of molecules in a relatively deep and spatially precise portion of posterior cortex, our EEG measurements were reflecting temporal variations in the activity of relatively superficial and widespread cortical neurons. Similarly, the causal determinants of oscillatory activities, and their variations over time, are not yet known with temporal or molecular precision; it may be the case that distinct cell types, neuromodulators, or long-range connections are influencing EEG signals which would not be reflected in the measurement of Glx and GABA+ in a single voxel within posterior cortices (Ahmad et al. 2022). The relations between EEG-based E/I ratios and MRS-based E/I ratio may also be nonlinear, such that different EEG analysis algorithms could capture different aspects of underlying MRS-based E/I ratio; larger datasets would be needed to determine whether an optimal combination of EEG-based measures could better account for MRS-based E/I ratio.

Our results also reinforce the fact that the phenomena we are considering may be varying over time on fundamentally different timescales. One of the most notable functions of variations in excitation and inhibition are their changes over developmental time, that is, years or even decades (Hensch, 2004; Takesian & Hensch, 2013), while in contrast, cognition-related modulation of E/I would necessarily operate on sub-minute timescales (Hellyer et al., 2016). The methods we considered provide clear tendencies toward longer or shorter timescales, with MRS-based measures needing a minimum of several minutes, NMM-based measures providing estimates which are nominally relevant on the sub-second level, and the other methods falling somewhere in between. It may be possible, then, that these measures simply do not reflect underlying cortical changes that operate on similar timescales, leading to a lack of evident within-person relations between them.

Robust links between EEG-based E/I ratio and MRS-based E/I ratio at a between-subjects level provides validation for EEG-based measures to be used in certain areas of research for which they have developed, namely, in investigating special populations such as individuals with ASD (Bruining et al. 2020), Alzheimer’s disease (van Nifterick et al., 2023), or monogenetic neurodevelopmental disorders (Geertjens et al., 2022). Our results indicate that the between-person differences in E/I revealed by contrasting these groups with healthy controls are likely to have common mechanistic bases (i.e. ratio of glutamate to GABA) whether they are measured using EEG or MRS. In contrast, for research which relies heavily on both within-person and between-person contrasts, such as the study of neuroplasticity (Shibata et al., 2017) or vigilance (Yang et al., 2023), EEG-based E/I ratio measures may not allow the assumption of mechanistic isomorphism.

While our results indicate that several recently developed algorithms for measuring E/I using EEG are indeed reflecting variations in the cortical ratio of glutamate to GABA, several limitations our present in our findings. The size of our dataset does not allow for precise estimates of effect sizes nor compelling rejection of candidate EEG-based E/I ratio measures, and the study of sleeping humans may lead to idiosyncratic results when contrasted to wakefulness; each of these limitations motivates future comparisons using concurrent multimodal imaging in awake humans.

## Acknowledgements

We would like to thank our funding sources: NIH (K99EY034891, R01EY019466, R01EY027841, R01EY031705, P30GM149405); NSF-BSF (BCS2241417). We would also like to thank Michael Freund for invaluable input when designing the statistical analyses.

## Supplemental Material

### Bayesian Multivariate Linear Mixed-Effects Model

The following is a schematic showing the specification of the Bayesian Multivariate Linear Mixed-Effects Model (BMVLMEM) estimated using **rstan** via **brms**, with each outcome variable modeled with a Gaussian response distribution. Within the same model, and estimated simultaneously, all outcome variables (to the left of the tilde) were predicted by a group-level intercept, an autoregressive term *ar()* of order 1 grouped by subject and with the number of the 5-minute block as the variable *time*, and a subject-level intercept. In addition, the correlation matrix of the subject-level intercepts was estimated (*subjectR*). Because this model specification accounted for effects of time and inter-individual differences, the remaining variance left unexplained was intra-individual variation. The correlation matrix of residuals, estimated in conjunction with the full BMVLMEM, therefore represented the within-subjects associations between the outcome variables. Note that our primary interest was linking each EEG measure with MRS-based E/I ratio, which we report in the main results. We also report the remainder of the correlation matrices, including correlations between EEG measures and with Glx and GABA+ estimates, below (see Figure S2). We tested the robustness of this model by fitting alternate models excluding the autoregressive terms as well as modeling the outcome variables as Student T distributions rather than Gaussian, but these did not change the qualitative pattern of results, and we report only the results of this primary model here.

**Figure S1.**
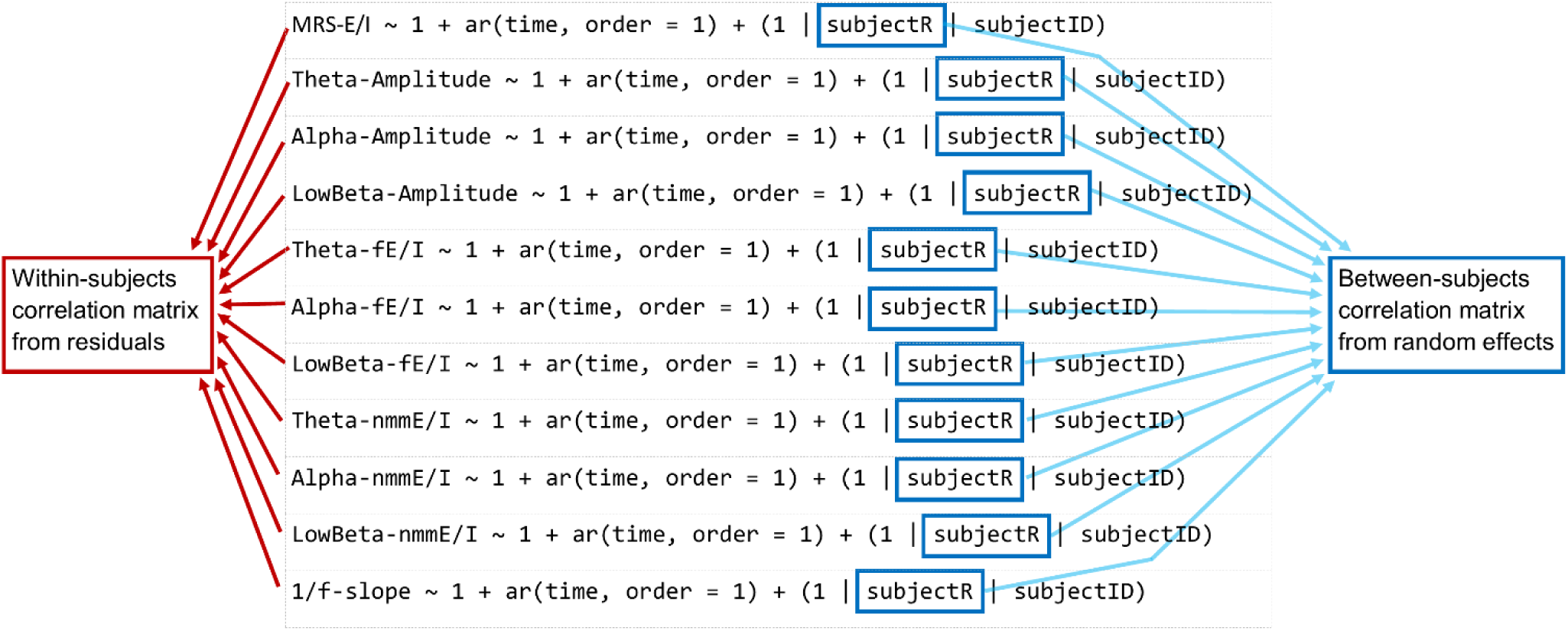
Bayesian Multivariate Mixed-Effects Model specification and correlation matrix estimation.

### Associations among EEG measures

In addition to our primary questions regarding links between EEG-E/I measures and MRS-based E/I, we additionally explored two additional questions. First, we assessed the extent to which different EEG-based measures of E/I were themselves collinear; second, we assessed the associations between each EEG-E/I measure was associated with either Glx or GABA+, as opposed to these molecules’ ratio in the form of MRS-E/I. To assess these relations we extracted the by-participant random intercepts and the residual correlations from the Bayesian multivariate linear mixed-effects model (see Figure S1 & S2; see also Figures 1b, 2b).

**Figure S2.**
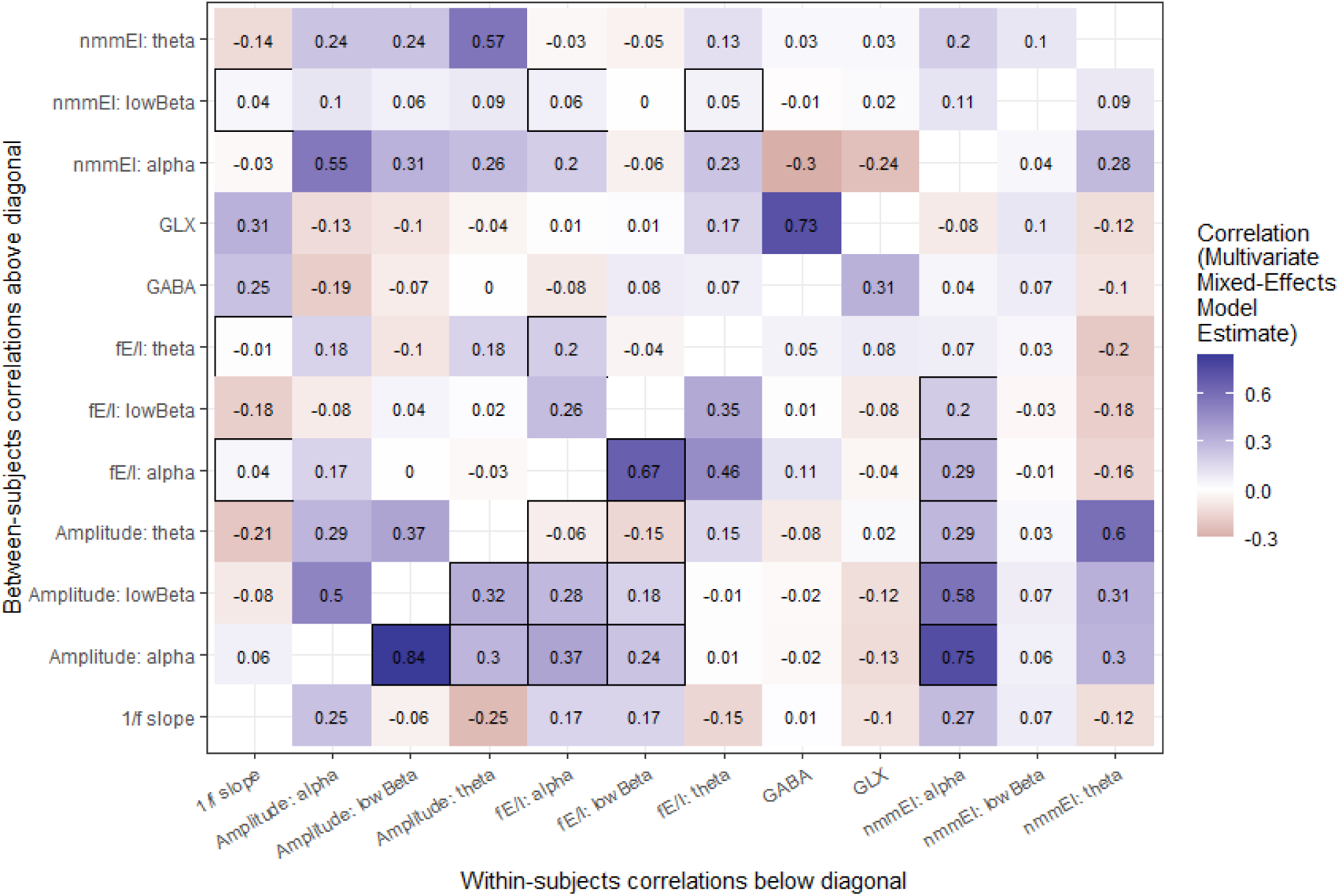
Correlations among measures apart from MRS-E/I (i.e., EEG measures as well as separated Glx and GABA+ concentration estimates). Above the diagonal are between-subjects correlations (see Figure 1b); below the diagonal are within-subjects correlations (see Figure 2b). Cells of EEG measures with reliable out-of-sample predictiveness of MRS-E/I are outlined. All estimates are drawn from a single multivariate mixed-effects model.

### Across-channel correlations

One way to assess the reliability of EEG-based measures, which were each estimated separately for each channel, was to examine the correlations between the same measure across the posterior channels we analyzed. Due to our interest in relatively macro-scale changes in E/I, as well as the basic properties of volume conduction (Burle et al. 2015), we expected that our EEG-based E/I estimates would have highly positive correlations with one another; this would indicate that the underlying algorithms are computationally stable and that our data quality was high. For most measures we do observe the expected large positive correlations between channels, although the NMM-based estimates are much more weakly inter-correlated (See Figure S3). Broadly, then, we see that measures of amplitude, fE/I, and 1/f slope all demonstrate expected cross-channel correlations indicating reliability, while the low correlations of NMM measures may be due to true underlying ability to measure independent measures from nearby electrodes, or may be due to computational instability of the NMM approach; further investigation of this topic may be of interest for future research.

**Figure S3.**
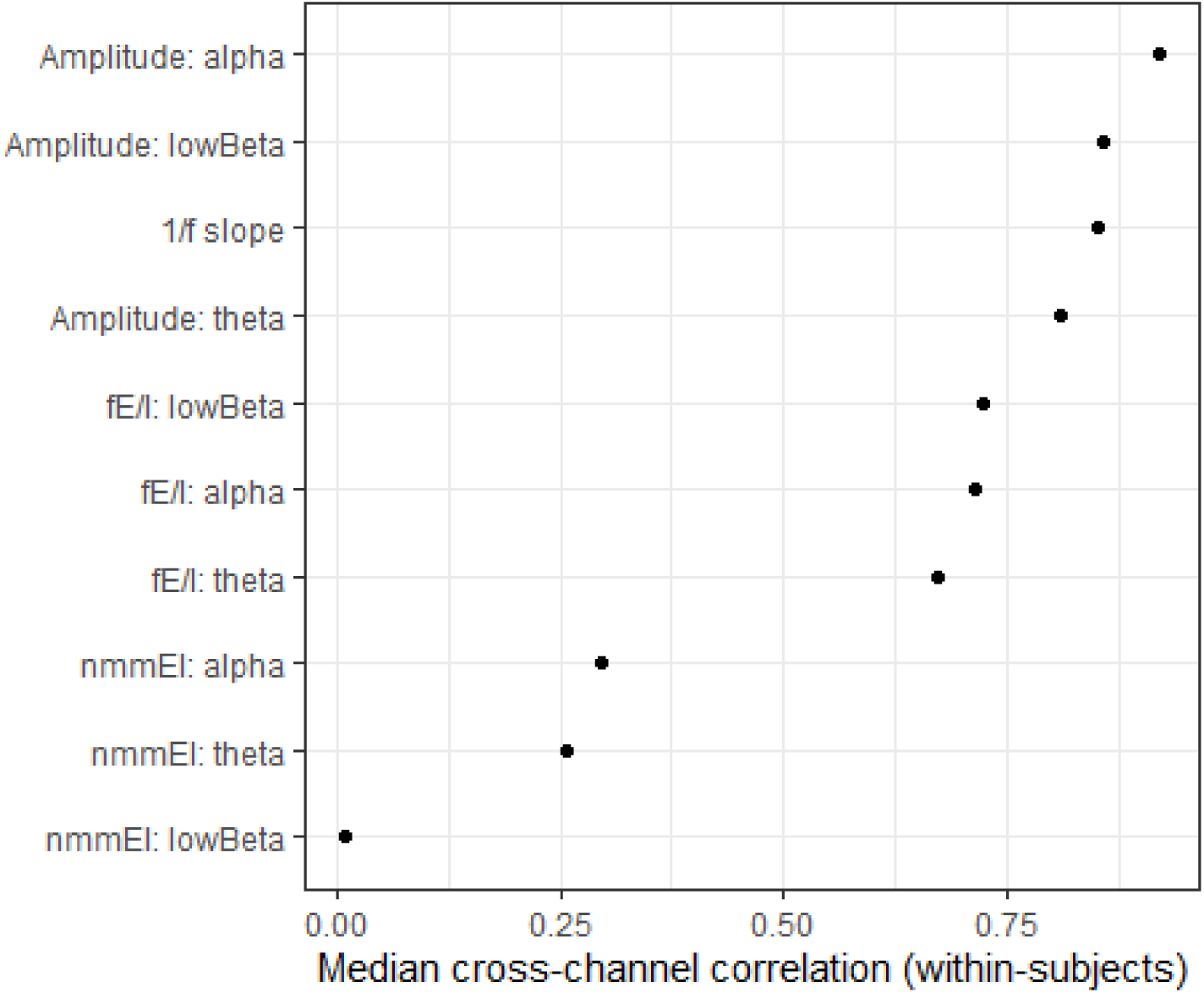
Median rank correlations between analyzed posterior channels’ estimates of EEG-E/I. Most estimates were highly correlated across channels, although NMM-based estimates had relatively low cross-channel reliability.

### Ratio of between-subjects to within-subjects variance

As a second exploration into the consistency of within-subjects estimates, which may influence the reliability of within-subjects E/I measures, we examined the ratio of between-subjects variance to within-subjects variance.

**Figure S4.**
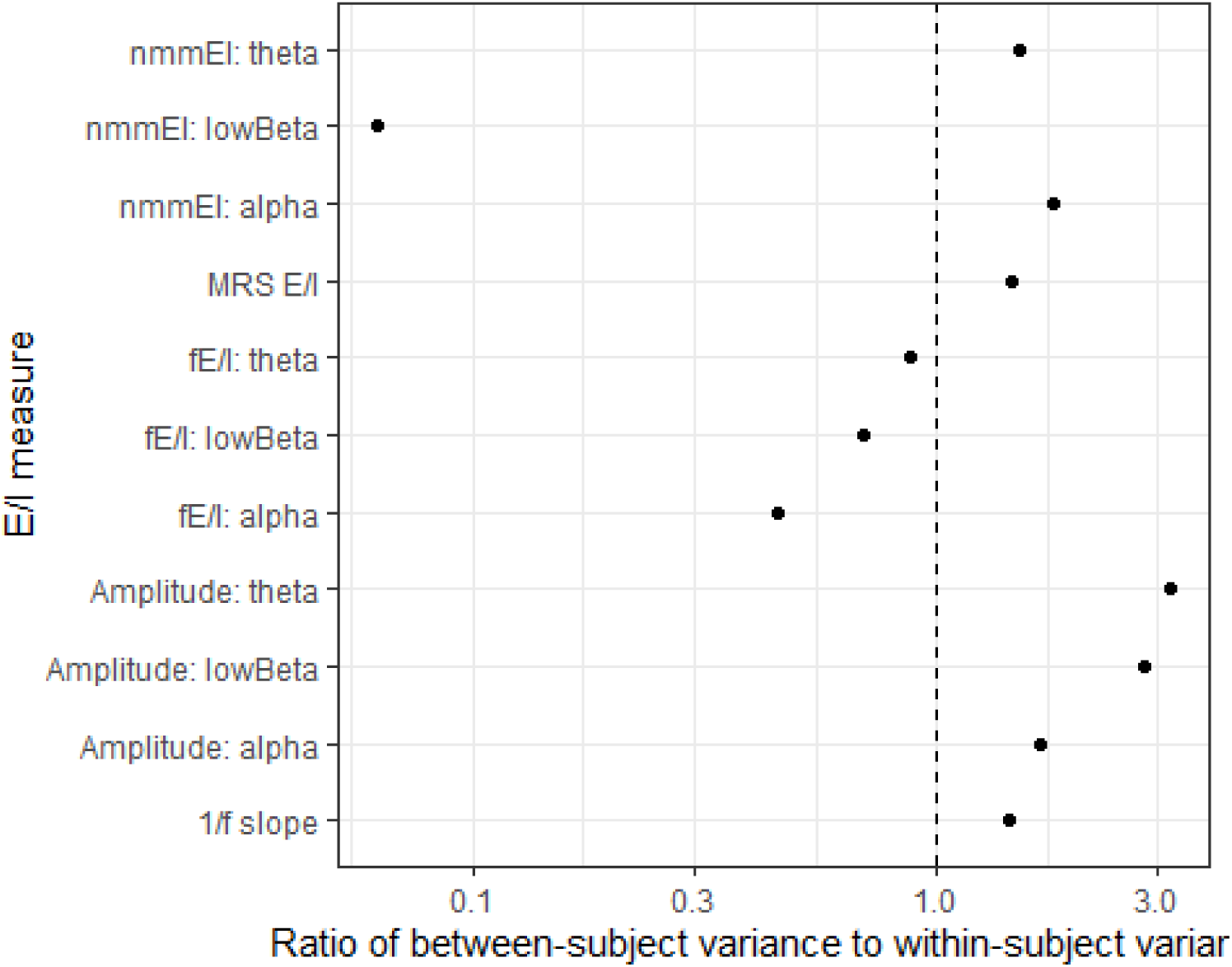
Ratios of between-subjects variances to within-subjects variances. Most measures had less than a 3:1 ratio of these variances, with MRS-based E/I having between-subjects variances approximately 1.45 times as large as within-subjects variances.

